# Effect of plant tubulin kinetic diversification on microtubule lengths

**DOI:** 10.1101/2021.05.11.443582

**Authors:** Kunalika Jain, Megha Roy, Chaitanya A. Athale

## Abstract

Microtubules (MTs) are dynamic polymers vital for cellular physiology. Bulk tubulin polymerization is nucleation dependent, while individual filaments exhibit ‘dynamic instability’ driven by GTP hydrolysis rates. Although MTs assembled from well-studied animal brain tubulins have very comparable nucleation and GTP-hydrolysis rates, the kinetic rates of evolutionarily more distant species could diverge. Here we focus on a plant tubulin, the legume *Vigna sp*. (mung bean) to test the effect of kinetic diversification on MT polymerization. We activity purify tubulin from seedlings and find MT filaments are fewer and shorter than animal brain tubulin. We find mung tubulin polymerization kinetics is nucleation dependent with a high rate of GTP hydrolysis and a critical concentration lower than previously reported for tubulins. A computational model of the kinetics based on the relative influence of rates of nucleation and hydrolysis demonstrates increased rates of hydrolysis can affect MT filament numbers and their lengths, as compared to increasing nucleation rates. Our approach provides a framework to compare the effect of evolutionary diversification of MT nucleation and elongation.

## Introduction

The role of microtubules (MTs) in cell growth and division has led to careful *in vitro* measurements of polymerization kinetics of tubulin. Theoretical models of microtubule polymerization can be categorised as either population dynamic models based on bulk kinetics or models single filament dynamics. Kinetic modeling approaches have been based on a variation of nucleation dependent polymerization (NDP) theory, based on which aggregation and polymerization occurs only above a threshold concentration, the critical concentration, c* (Ferrone et al., 1985, Bishop and Ferrone, 1984). Kinetic measurements and modeling demonstrating GTP-hydrolysis in the lattice and differential rates of tubulin addition depending on whether GTP or GTP bound, demonstrated that a GTP-tubulin gradient would emerge from the growing end, the GTP-cap (Carlier and Pantaloni, 1981). A model of MTs with two dynamic ends growing in solution with a GTP-cap at one end (Hill, 1985) could reproduce experimental MT length distributions from filament shearing experiments (Chen and Hill, 1985). The single filament ‘dynamic instability’ of MTs transitioning from growing to shrinking statescatastrophe- and the reverse transitionrescue (Mitchison and Kirschner, 1984) were captured in model with four parameters-two transition frequencies and the rates of growth and shrinkage (Dogterom and Leibler, 1993). The model could explain the regulation of the frequency of catastrophe when cells transitioned from ‘unbounded’ growth in interphase to ‘bounded’ growth with rapid dynamic in mitosis (Verde et al., 1992). The loss of the GTP cap arising from the hydrolysis of GTP-tubulin were shown to result in a transition to shrinking MTs (Dogterom and Leibler, 1993, Flyvbjerg, Holy and Leibler, 1996, Flyvbjerg et al., 1994). A detailed GTP cap with protofilament dynamics could explain the delay between tubulin monomer dilution and catastrophe (Brun et al., 2009) and describe the expected shape of a single filament tip (Gardner et al., 2014). A recent model has proposed that hydrolysis rates determine the separation of two critical concentrations of elongation and nucleation (Jonasson et al., 2020). A test of such models requires a comparison with experimental data with varying hydrolysis rates with most studies focussing on the kinetics of animal microtubule polymerization.

Plant tubulins have been previously isolated from multiple sources with some of the earliest reports describing tubulin isolation from ‘mung’ (Azuki) bean (*Vigna radiata*) (Mizuno et al., 1981). A comparison of filament assembly and drug-binding sensitivity of tubulins isolated from animal brain and cultured cells of multiple higher plants, suggested plant tubulins bind less effectively to colchicine as compared to animal tubulins (Morejohn and Fosket, 1982). These differences in sensitivity to drug binding could be explained by the divergence of *α* tubulin sequences in plants from animals, as compared to the more conserved *β*-tubulin sequences (Morejohn and Fosket, 1984, Morejohn et al., 1984). MT filaments from a diverse set of plant tubulins-mung bean, pea, zucchini, cucumber seedlings and carrot cell suspensions-were shown to require only GTP, Mg^2+^ ions, EGTA and a crowdant for assembly (Mizuno, 1985). More recently the development of a TOG-affinity based tubulin isolation method (Widlund et al., 2012) has been successfully employed to isolate and kinetically characterize tubulin from multiple plant sources (Hotta et al., 2016). The tubulin filaments from *Arabidopsis* were found to be more dynamic and shorter than animal tubulins. The observed differences between plant and animal tubulins in terms of polymerization kinetics and filament length distributions have however not yet been compared using a theoretical framework for a comprehensive picture of evolutionary diversification.

Here, we optimize a method to activity isolate tubulin from (*Vigna sp*.) (mung bean) seedlings and compare their polymerization kinetics to better studied animal brain tubulins. We measure filament length distributions in label free microscopy and estimate polymerization rates from turbidimetry kinetics. Rescaling analysis is used to compare the nucleation dependent polymerization with goat and porcine brain tubulin. We find mung MT polymerization kinetics diverge from animal brain tubulin both in terms of GTP hydrolysis rates and the critical concentration. We develop a computational model to examine whether the rapid kinetics of nucleation or GTP hydrolysis dominate MT polymerization.

## Results and Discussion

### Comparing microtubule filament length distributions of plant and animal tubulin

Seedlings are expected to be enriched for tubulin due to rapid growth and have therefore been successfully used in the past as a source for tubulin isolation, such as from the legume *Vigna sp*., mung bean (Mizuno et al., 1981, 1985, Sen et al., 1987). We proceeded to produce rapidly growing mung seedlings as a source of tubulin (Figure 1(A)) and optimized the temperature-dependent polymerization and depolymerization method used to isolate brain tubulin reported previously (Castoldi and Popov, 2003) to minimize the presence of MAPS. The prominent 50 kDa band in the sample (Figure 1(B)) was confirmed to be tubulin by immunostaining dot blots with an anti-*Arabidopsis α*-tubulin antibody (Figure 1 S1(a)). Interestingly, the same antibody also recognizes goat and porcine brain tubulin, but not BSA, suggesting the epitopes recognized in plants are also conserved in animals, consistent with phylogenetic distances between tubulins (Figure S1(b)). We assembled filaments of mung and goat tubulin in the presence of GMPCPP, the non-hydrolysable analog of GTP. Microscopic images in label free interference reflection microscopy (IRM) suggest mung MTs are shorter than goat brain MTs (Figure 1(C,D)). The filament length distribution of goat MTs with a mean length of 1.57 *µ*m, is similar to the mean length of mung MTs of 0.98 *µ*m. However, goat MT lengths have more outliers as seen from the fit to a lognormal distribution (Figure 1(E)). We proceeded to ask whether the qualitatively similar filament length distribution with small, quantitative differences, could be explained by a similarity in the kinetics of polymerization.

**Figure 1:**
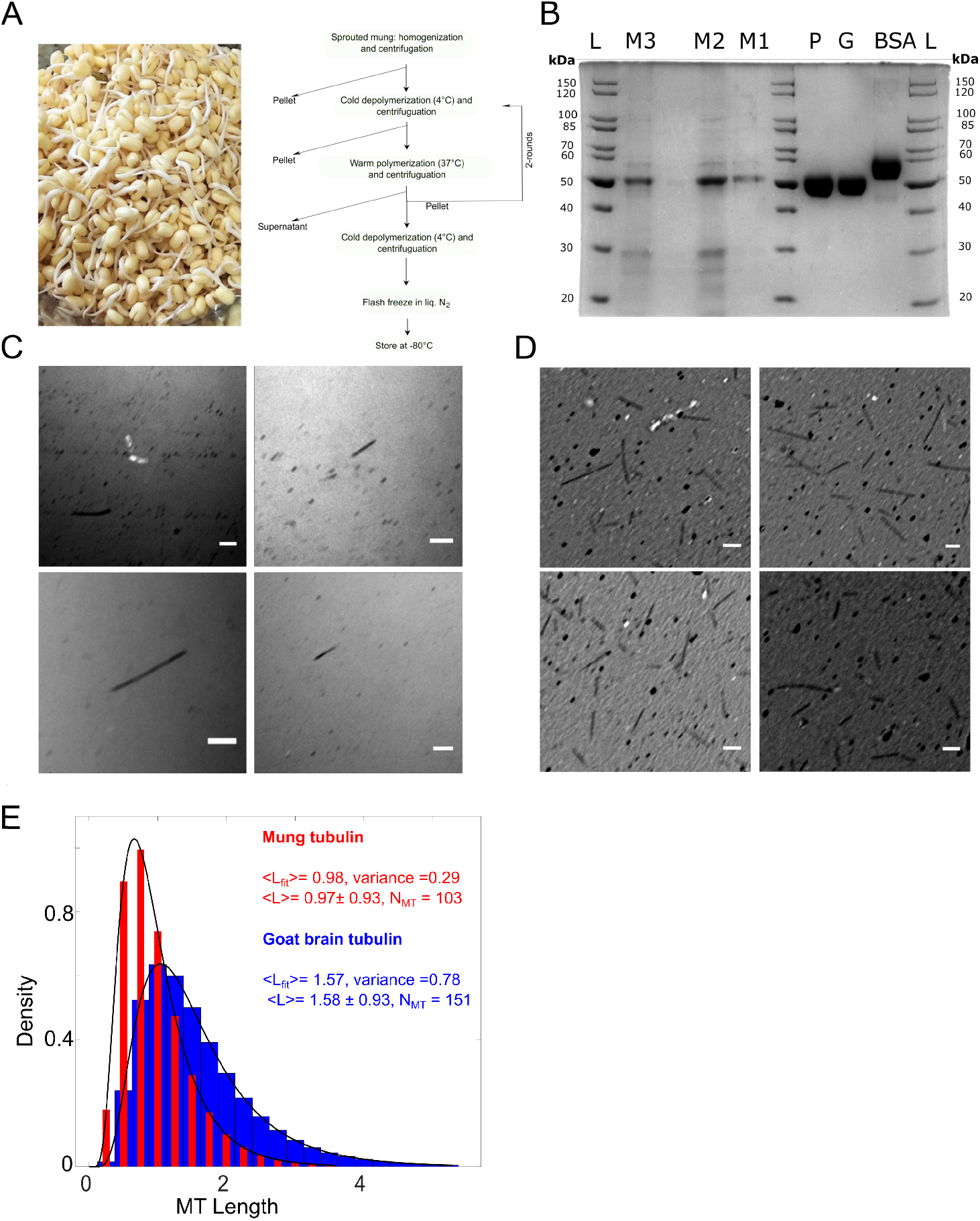
Comparison of microtubule (MT) filaments assembled from mung and goat brain tubulin. (A) Sprouted beans of *Vigna radiata* (mung) were used to isolate tubulin using the temperature dependent polymerization-depolymerization method. (B) The multiple mung isolates (M1, M2, M3) separated on a 10% SDS-PAGE gel stained with Comassie brilliant blue (CBB) have a prominent 50 kDa band comparable in size to P(orcine) and G(oat) tubulin. BSA is a control. L: Marker. (C-D) IRM images of multiple regions of interests (ROIs) MT_s_ assembled from (C) 0.75 uM mung tubulin and (D) 30 μM goat brain tubulin in BRB-80 with 1 mM GMPCPP incubated for 1 hour at 37°C are seen. Scale: 1 *μ*m. (E) The frequency distribution of filament lengths of MTs assembled from mung (red, 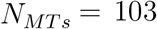) and goat brain tubulin (blue, 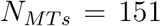) are plotted and fit to a lognormal distribution. < *L_fit_*>: mean length from fit, L: arithmetic mean.

### Polymerization kinetics to measure nucleation dependence and critical concentration

The polymerization kinetics of plant tubulin isolated from *Arabidopsis* cell suspension cultures were reported to be rapid at single filament level with a critical concentration c* of 7.5 *µ*M (Hotta et al., 2016). We find mung tubulin polymerization in presence of GTP resulted in a transient increase and rapid fall in absorbance (Figure 2(A)), comparable to the rapid ATP hydrolysis driven kinetics of ParM filaments (Garner et al., 2004). The critical role of GTP concentration relative to tubulin is seen when GTP was added 10 minutes after incubation demonstrating increased polymerization, only when the GTP:tubulin concentration ratio was sufficiently high (Figure S2). At the same time, stabilization by paclitaxel (referred to as taxol) resulted in slower kinetics and three phases of polymerization became visible-lag, elongation and saturation (Figure 2 S3). The GTP kinetic curves were fit to a saturation model of polymerization kinetics (Equation 1) to estimate the polymerization rate, *r* (1/min), characteristic time *τ* (min), and maximal polymer mass *A*_*max*_ (arbitrary units) for increasing concentrations of tubulin from mung (Figure 2(A)), goat brain Figure 2(B)) and commercial porcine brain tubulin Figure 2(C)).

**Figure 2:**
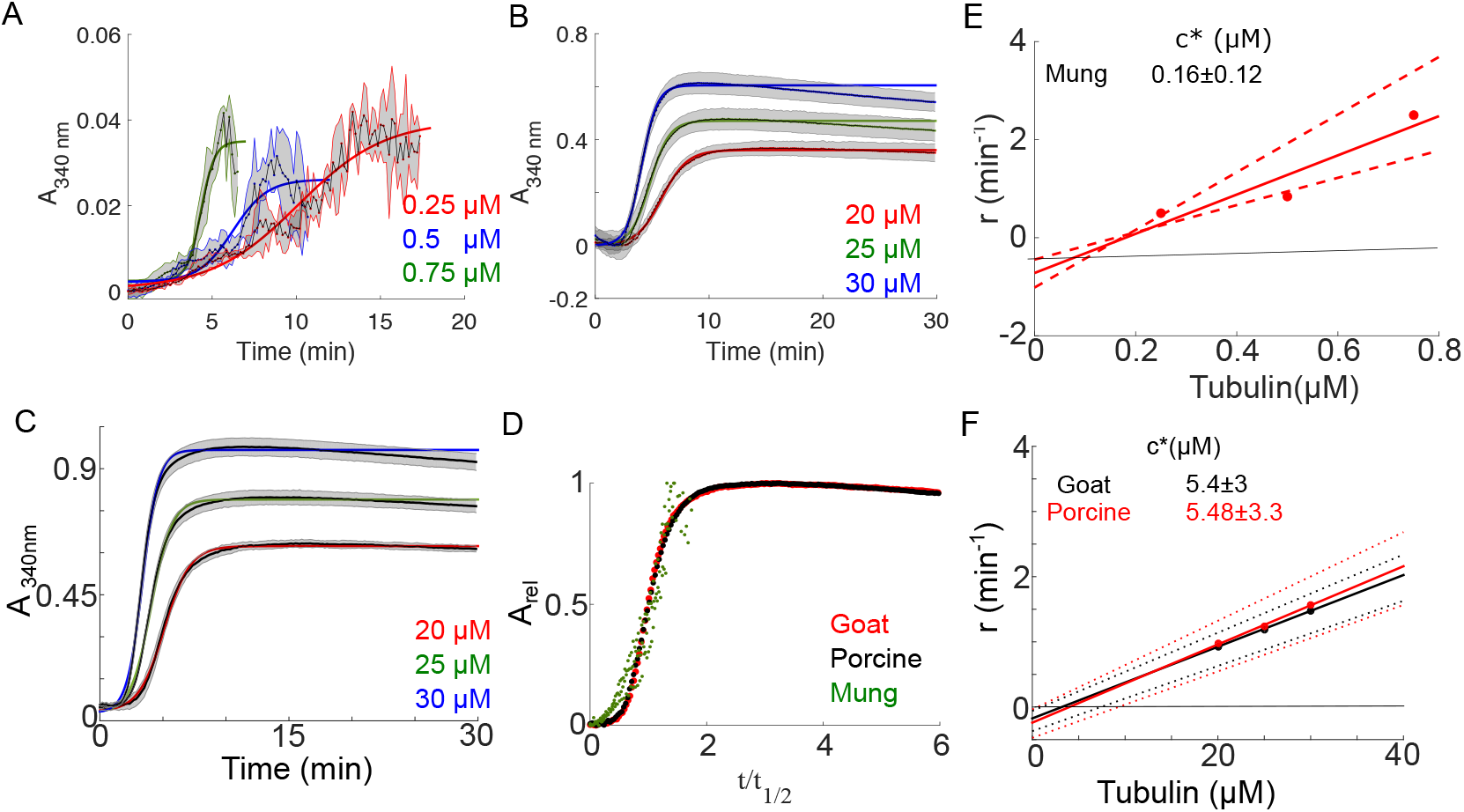
Comparative analysis of tubulin polymerization kinetics from plant and animal sources. (A) The polymerization kinetics of 0.25 to 0.75 *µ*M mung tubulin were measured by the absorbance at 340 nm over time in presence of 10 mM GTP. Mean values (black lines) were fit to a saturation model (Equation 1) to estimate the polymerization rate (r), half time (t_1/2_), and maximal absorbance (A_*max*_). Grey areas indicates s.d. (n=3). (B,C) Similar absorbance based polymerization kinetic curves of (B) goat and (C) porcine brain tubulin (20 to 30 *µ*M) were measured in presence of GTP and mean curves (black lines) and fit (coloured lines). (D) The kinetic data was rescaled as *A*_*rel*_ = (*A*(*t*) *-A*_0_)/(*A*_*max*_ *-A*_0_) and t/t_1/2_, as described in the Methods section. Filled circles are individual data points for multiple concentrations of mung (green), goat (red) and porcine tubulin (black). (E,F) The polymerization rate *r* (per minute) of tubulin as a function of tubulin concentration (circles) is fit by a linear model (solid line). The critical concentration inferred from the x-intercept of the line is (E) 0.16 *µ*M for mung tubulin and (F) 5.4 *µ*M for goat brain (black) and 5.5 for porcine brain tubulin (red). The dashed-lines indicate the uncertainty from the 95 % confidence interval.

Rescaling the diverse tubulin polymerization kinetics as described before (Flyvbjerg, Jobs and Leibler, 1996) resulted in the mung tubulin data falling on the same curve as goat and porcine brain tubulin (Figure 2(D)). This suggests that mung tubulin polymerization is nucleation dependent, similar to animal brain tubulin. The critical concentration of mung tubulin was estimated to be 0.16 *µ*M (Figure 2E), while brain tubulin from goat and porcine sources have c* of *∼* 5 *µ*M (Figure 2F), based on the increasing polymerization rate *r* as a function of tubulin concentration. The animal tubulin values are comparable to previous reports (Bonfils et al., 2007), while the trend of plant tubulin c* being lower than animal is consistent with previous reports (Table S1). Our report is however the lowest reported value for plant tubulin so far.

MTs are nucleated in animal somatic cells *in vivo* primarily by centrosomes that form radial arrays or asters (Karsenti et al., 1984). Such nucleators function by lowering the critical concentration for nucleation, c* (Buendia et al., 1992). Despite the fact that plant cells lack centrosomes, well organized MT arrays are formed in interphase and mitosis. In such cells, the function of centrosomes appears to be replaced by three pathways as reviewed by Yi and Goshima (Yi and Goshima, 2018): (a) nucleation from small microtubule organizing centers (MTOCs), (b) MT-dependent MT nucleation and (c) altered minus-end dynamics due to either intrinsic properties of tubulin or by the action of MT-associated proteins (MAPs). A sub-micromolar critical concentration is therefore lower by an order of magnitude to typical cellular tubulin concentrations of a few micromolar (Loiodice et al., 2019), suggesting an additional mechanism by which plants might overcome the absence of discrete nucleation centres, intrinsic to tubulin.

Thus while mung tubulin kinetics are dramatically different from animal brain tubulin-nucleation at a lower concentration and faster hydrolysis-experimental data suggests filament lengths are very similar. In order to explain this apparent paradox, we simulate microtubule polymerization dynamics.

### Stochastic model of nucleation and hydrolysis of a population of MT filaments

We have developed a model of MT nucleation and growth dynamics using a stochastic simulation engine of cytoskeletal mechanics, Cytosim. In the simulation, multiple MTs can be nucleated, elongate and shrink stochastically in a 2D geometry, surrounded by a pool of GTP-tubulin monomers (Figure 3(A)). Polymerization of MTs begins with the aggregation of monomers into oligomers determined by a nucleation rate *r*_*nuc*_. A lower critical concentration of nucleation implies a more rapid nucleation rate. Filaments elongate with an effective velocity of growth *v*_*g*_, by the addition of GTP-bound tubulin dimers to the plus-end. Within each filament the GTP is hydrolyzed by tubulin to GDP in a vectorial manner at a rate *r*_*hyd*_, from the minus to the plus-end. If a filament has only GDP bound monomers at its plus-tip, it undergoes a transition from growing to shrinking state. The transition is referred to as a catastrophe, and the filament length reduces at a rate determined by the shrinkage velocity, *v*_*s*_. Since MTs can only grow by end-on addition of monomers, a high rate of nucleation is expected to result in many ends being formed and result in a kinetic competition between new oligomer formation and the elongation of the oligomers. MT stability is determined by the opposing tendencies of filament growth by addition of GTP-tubulin and the GTP hydrolysis rate, that combine to determine the GTP-cap size (Roostalu et al., 2020). We examine two scenarios, (a) *nucleation-based* in which nucleation rates *r*_*nuc*_ are increased for a low hydrolysis rate and (b) *hydrolysis-based* in which the GTP hydrolysis rate *r*_*hyd*_ increases for a fixed nucleation rate.

**Figure 3:**
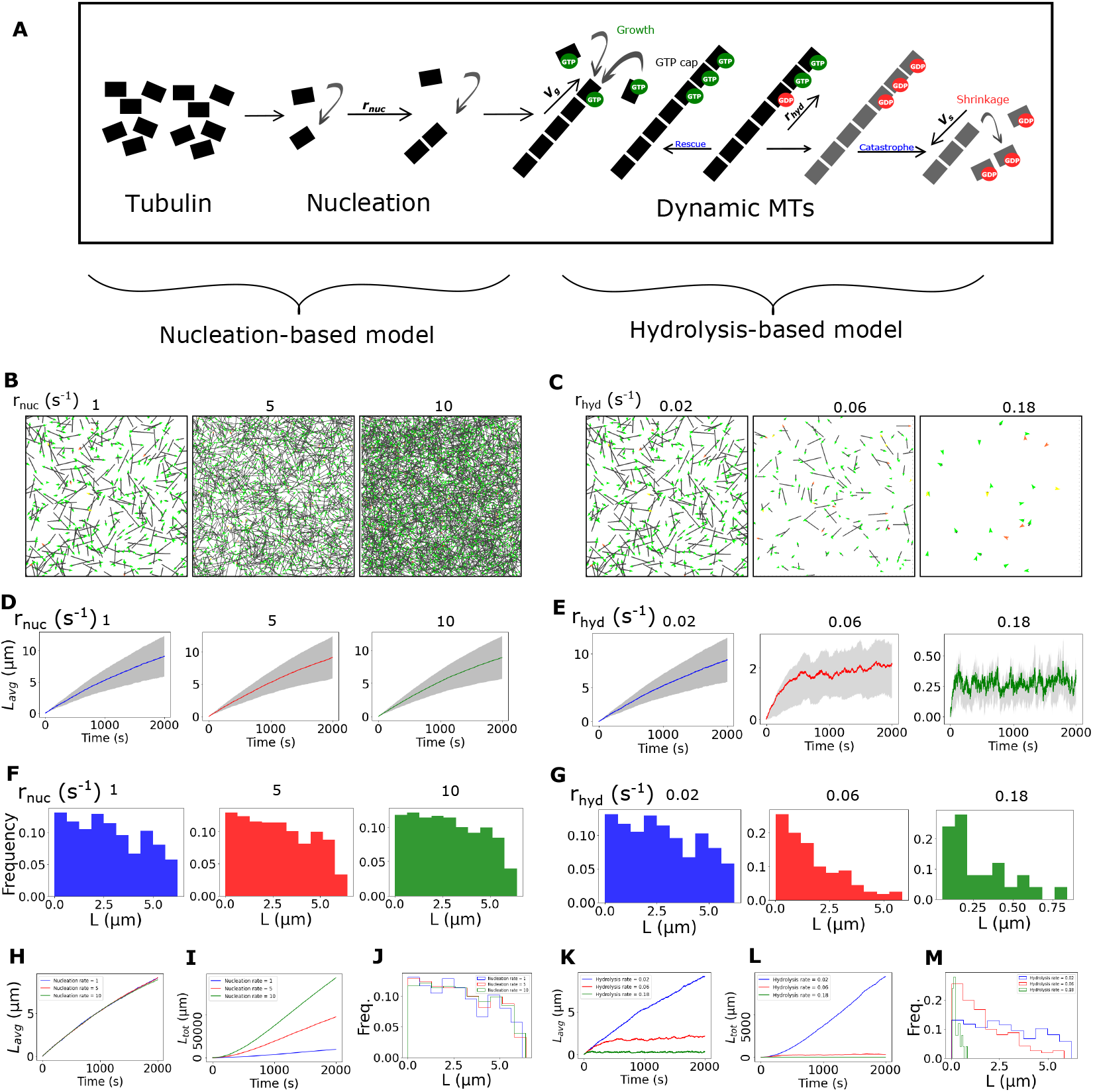
Simulating the kinetic competition between MT nucleation and GTP-hydrolysis. (A) The schematic depicts the model of tubulin monomer nucleation, oligomerization and filament dynamics as microtubules, with characteristic growing speed shrinking speed and hydrolysis rate. Model parameters are taken from literature (Table 1). (B, C) We compare simulations of filament distribution when either (B) the nucleation rate (*r*_*nuc*_) was varied while the hydrolysis rate *r*_*hyd*_ was 0.02 *s*^-1^ or (C) *r*_*hyd*_ was varied for a constant nucleation rate of 1 *s*^-1^. Simulation snapshots are shown for 500 sec. (D,E) The time-dependence of mean length of the population (coloured line) with time is plotted with the s.d. (grey area). (F,G) The frequency distribution of MT lengths at the end of simulation time is compared. (H,K) The mean length with time, (I,L) total MT mass with time and (J,M) frequency distribution of lengths at the end of simulation time are compared for the respective nucleation rate or hydrolysis rate variation.

### Simulating the effect of kinetic competition between nucleation and hydrolysis on filament lengths

In simulations we examined the effect of increasing the nucleation and hydrolysis rates, for a monomer pool limited to 7.5 *µ*M and kinetic parameters either taken from those reported for *Arabidopsis* MT dynamics or varied in the absence of available values (Table 1). We observe increasing the nucleation rate results in the increased number of filaments (Figure 3(B), SV1(a)), while an increasing hydrolysis rate result in shorter, more short lived and fewer filaments (Figure 3(C), SV1(b)). This matches intuitive expectations that if there is more nucleation more ends will be formed, while if hydrolysis is greater, the filaments that form will be more unstable, undergoing catastrophes and becoming both fewer and shorter. The quantitative analysis of the simulations shows that while the mean length (*L*_*avg*_) constantly increases with *r*_*nuc*_ for the *nucleation-based* model (Figure 3(D)), it rapidly reaches a steady state for the *hyrolysis-based* model (Figure 3(E)). The MT length distribution in the *nucleation based* model appears uniform at any given time (Figure 3(F)), while it appears to follow an exponential decay at high values of *r*_*hyd*_ in the *hydrolysis-based* model (Figure 3(G)). Interestingly, when we increase the nucleation rate, corresponding to reducing critical concentration, *L*_*avg*_ remains unaffected (Figure 3(H)). We interpret this in terms of the fact that the total MT length (a surrogate for total polymerized tubulin concentration) never exceeds the maximal liming pool (Figure 3(I)). The distributions of length are also constant for the values of *r*_*nuc*_ tested (Figure 3(J)). In contrast the *hydrolysis-based* model predicts varying *r*_*hyd*_ results in a clear changes in mean MT lengths (Figure 3(K)), total MT length (Figure 3(L)) and result MT length distributions (Figure 3(M)).

**Table 1:** The simulation parameters used model nucleation and polymerization kinetics of MT filaments.

| Parameters. | Value | Units | Reference |
| --- | --- | --- | --- |
| **MT polymerization dynamics** |  |  |  |
| Nucleation rate | 1 to 10 | $s^{-1}$ | This study |
| Hydrolysis rate | 0.02 to 0.18 | $s^{-1}$ | This study |
| Growth velocity ($v_g$) | 0.012 | $\mu\text{m}\cdot s^{-1}$ | (Hotta et al., 2016) |
| Shrinkage velocity ($v_s$) | 0.826 | $\mu\text{m}\cdot s^{-1}$ | (Hotta et al., 2016) |
| **Simulation setup** |  |  |  |
| Time step | 0.02 | s | This study |
| Filament section | 0.5 | $\mu\text{m}$ | (Khetan and Athale, 2016) |
| Simulation box (periodic) | $40\times 40$ | $\mu\text{m}\times\mu\text{m}$ | This study |
| Total simulated time | 2000 | s | This study |
| Viscosity ($\eta$) | 0.9 | $\text{mPa}\cdot\text{s}$ | Water |
| Thermal energy (kBT) | 4.2 | pN nm | - |
| Total polymer mass | $7.5\ \mu\text{M}$ | Molar concentration | (Hotta et al., 2016) |

The comparison between predictions of the *nucleation-based* and *hydrolysis-based* models predicts that MT lengths are not affected by order of magnitude changes in the nucleation rate, but comparable changes in the hydrolysis rates do alter lengths. This would suggest that the divergence of critical concentration and hydrolysis rates both seen between mung and animal tubulin and the resulting change in length distributions, can be best explained by a *hydrolysis based* model.

*In vivo* MT organization in plant cells is distinct from that seen in animal cells, with plant MTs forming cortical arrays (Allard et al., 2010, Elliott and Shaw, 2018, de Keijzer et al., 2014) that regulate cell wall synthesis (Oda, 2015), while in animal cells MTs typically form tracks for transport in interphase and spindles in mitosis. Taken together plant MT lengths are expected to be shorter than those of animal MTs. Theoretical models of plant MT length distributions have described the role of severing proteins in affecting the qualitative nature of MT length distributions (Tindemans and Mulder, 2010), considered important for the cortical aligment of plant MTs at interphase (Deinum et al., 2017). The lengths of filaments in our samples are however unlikely to be the result of severing by katanins, since it also requires a dense MT network of MTs for its activity (Fan et al., 2018). The simple model of a kinetic competition between nucleation and GTP-hydrolysis invoked here to explain the observed *in vitro* MT length distributions, is supported by evidence from single-filament dynamics of MT filaments whose lengths and numbers were increased by the addition of a E254A mutant *α*-tubulin that had a reduced rate of GTP-hydrolysis (Roostalu et al., 2020). It would be interesting to see whether additional MAPS in the plant tissue regulate the rate of GTP-hydrolysis *in vivo*, and consequently the number and lengths of MTs.

## Conclusions

In summary, we find the activity isolated plant tubulin from the mung bean (*Vigna sp*.) forms MT filaments whose average lengths are similar to that of goat brain MTs. At the same time kinetics of GTP hydrolysis and nucleation are much faster, with the critical concentration of mung tubulin *∼*30 times smaller than brain tubulin. To resolve this apparent contradiction, we tested a computational model that includes tubulin nucleation and GTP-cap. We show higher hydrolysis rates rather than nucleation rates, can better account for the small quantitative differences in MT length distributions of tubulins with divergent kinetics.

## Materials and methods

### Activity-based purification of tubulin

Tubulin was purified from germinated seedlings of *Vigna radiata* and brains of freshly sacrificed goats. Both purifications involved a temperature based polymerization-depolymerization cycle as described previously (Castoldi and Popov, 2003), as described briefly in the following text. Mung seeds were germinated at 30^*o*^C in a commercial ‘sprouter’, a 2-chamber container to separately soak and sprout, resulting in seedlings after 12 hour of soaking and 36 hours of germination. The manually de-husked seeds were homogenized in a heavy duty Waring blender (Stamford, CT, USA) and the homogenate clarified by high speed centrifugation in a Sorvall Ultracentrifuge (Thermo Scientific, Waltham, MA, USA) at 10^5^ g at 4°C for 40 min. The supernatant was processed for subsequent polymerization and depolymerization cycles. Polymerization was achieved by incubating the lysate with 1 M PIPES, 33% glycerol, 1.5 mM ATP and 0.5 M GTP at 37°C for 45 min. The polymerized sample was pelleted by centrifugation at 150,000 g at 32°C for 45 min. Depolymerized was achieved by the addition of cold BRB80 buffer (80 mM Pipes, 1 mM EGTA, 1 mM MgCl_2_) supplemented with 0.1 M NaCl by incubation at 4°C for 30 min. The sample was then centrifuged at 76,000 g for 30 min to retain the depolymerized supernatant. This cycle was repeated and the final pellet was depolymerized and resuspended in BRB 80 buffer. For comparison, tubulin from goat brains was similarly extracted by using freshly sacrificed animals. Brains were cleaned to remove blood clots, homogenized, centrifuged to clarify and the same polymerization-depolymerization cycles in high-molarity PIPES buffer were followed as before. Unlabelled porcine tubulin was obtained commercially (Cytoskeleton Inc., USA). All fine chemicals were sourced from Sigma-Aldrich, Mumbai, India, unless otherwise stated.

### Immunoblotting

Dot blots were made by pipetting mung, goat and porcine tubulin samples on a nitrocellulose membrane (Thermo Fisher, USA). The membrane was blocked with 5% milk powder in TBS-T buffer (w/v) for 1 hour at room temperature, followed by 1 hour incubation at 4 °C with a 1:5000 diluted polyclonal rabbit anti-Arabidopsis *α*-tubulin antibody (AS10 680, Agrisera, Sweden) primary antibody (primary). Membranes washed in TBS-T buffer (4 washes, each for 15 mins) were incubated with HRP-conjugated secondary antibodies at a dilution of 1:10,000 for 1 hour at room temperature and developed with a chemiluminescent reagent. HRP-conjugated goat anti-rabbit IgG (H&L) (AS09 602 antibody, Agrisera, Sweden) secondary antibody was developed using ECL SuperBright Reagents (Agrisera, Sweden). Blots were imaged using a chemiluminescence imaging system (G-Box, Syngene, Frederick, MD, USA).

### Filament microscopy

MT filaments were prepared using unlabelled tubulin from mung seedlings and goat brains. MTs were prepared either with 0.75 *µ*M mung tubulin or 30 *µ*M goat brain tubulin in BRB-80 buffer with 1 mM GMPCPP (Jena Bioscience, Germany) and incubated at 37°C for 60 min. The filaments were flowed in chambers made by sandwiching a 100 *µ*m thick double backed tape (Ted Pella Inc., CA, USA) between a glass slide and coverslips #1.5 of 0.17 mm thickness (VWR, Avantor, USA). Slides and coverslips were acid washed in 0.1 M HCl for 2 hours at room temperature followed by several washes of Milli-Q water to remove any traces of acid. Microscopic images were acquired using 100x NA 0.95 oil lens on a Nikon TiE (Nikon Corp., Japan) inverted epifluorescence microscope with a cooled CCD camera Andor Clara2 (Andor, Oxford Instruments, U.K.). A lexan stage enclosure with a temperature control system (Oko Labs, NA, Italy) was used to maintain temperature at 30°C. Interference reflection microscopy (IRM) images were acquired with a 50/50 beamsplitter (Chroma Technology Corp, VT, USA) in the reflected light path based on a previously described method (Mahamdeh and Howard, 2019, Mahamdeh et al., 2018).

### Fitting and rescaling of tubulin polymerization kinetics

Tubulin polymerization kinetics were measured for concentrations ranging betwee 0.25 *µ*M and 2 *µ*M were polymerized by incubation with BRB-80 buffer containing 10% glycerol and either 1 mM GTP (Jena Bioscience, Germany) with 1 mM MgCl_2_ or 10 *µ*M taxol, paclitaxel (Cytoskeleton Inc., CO, USA) with 5 mM MgCl_2_. Kinetics of tubulin were measured by turbidimetry by following absorbance at 340 nm every 10 s for a period of 30 min using a plate reader (Varioskan, Thermo Scientific, USA) with samples placed in well UV-transparent half-area flat-bottomed plates (Sigma Aldrich, Mumbai, India). In experiments where GTP was initially left out and then added 10 min after the reaction had proceeded the absorbances were path length corrected to account for changes in volume. Similarly, 15 to 30 *µ*M of goat brain tubulin isolated in lab, or porcine brain tubulin (Cytoskeleton Inc., CO, USA) were incubated with 1 mM GTP, 60% glycerol and 5 mM MgCl_2_ at 37°C and kinetics measured similar to mung tubulin. The time-dependent absorbance (*A*_*t*_) kinetics were fit to a standard four-parameter model used previously to fit for nucleation limited polymerization kinetics (Nielsen et al., 2001, Sabareesan and Udgaonkar, 2014, Schummel et al., 2017) which takes the form:

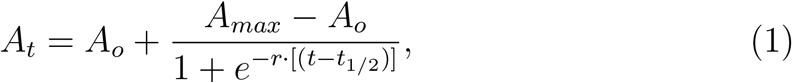

where *A*_*o*_ is the initial polymer mass, *A*_*max*_ is the maximum polymer mass, *t* is time over which the kinetics are measured in minutes, *t*_1/2_ is the half-maximal time of polymerization kinetics, and *r* is the polymerization rate per minute (Nielsen et al., 2001, Schummel et al., 2017). This polymerization rate *r* = 1/*τ*, where *τ* is the characteristic time constant described previously (Sabareesan and Udgaonkar, 2014).

Kinetic data was rescaled in order test the nucleation limitation of tubulin polymerization kinetics based on the approach described by (Flyvbjerg, Jobs and Leibler, 1996) as follows: (a) the time-dependent absorbance *A*_*t*_ was normalized to relative absorbance *A*_*rel*_, (b) the half-maximal time of the kinetics *t*_1/2_ was estimated, and (c) time was rescale by dividing it by *t*_1/2_. Relative absorbance was estimated by scaling for the difference between the initial *A*_0_ and maximal values *A*_*max*_, using the expression:

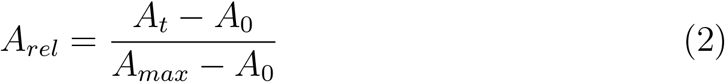

In addition to this scaling of the absorbance, a second scaling is performed for the time based on previous work (Flyvbjerg, Jobs and Leibler, 1996, Sabareesan and Udgaonkar, 2014), to estimate the half-maximal time *t*_1/2_. This estimate was obtained by fitting individual absorbance curves (*A*) to the four parameter kinetic model of Equation 1.

### Simulation

The model was developed in Cytosim (https://gitlab.com/f.nedelec/cytosim/ cloned in Feb 2021), a general C++ simulator of cytoskeletal dynamics and motor mechanics (Nedelec and Foethke, 2007). Filaments were nucleated at a constant nucleation rate (*r*_*nuc*_) and positioned throughout space randomly. The kinetic model was based on a velocity of growth (*v*_*g*_) that determines the rate of addition of GTP bound monomers, shrinkage velocity (*v*_*s*_) and the vectorial hydrolysis rate (*r*_*hyd*_) that results in the conversion of GTP to GDP bound tubulin. The parameters of the model were taken from experimental values reported for *Arabidopsis* tubulin dynamics (Hotta et al., 2016) whenever available, and scanned for those that were not reported previously.

## Acknowledgments

We are grateful to Mohammed Mahamdeh and Kheya Sengupta for help setting up interference reflection microscopy, to Ranjith Padinhateeri for discussions about MT dilution experiments and Uma Shaanker for discussions about plant tubulins. Kaveri Vaidya and Tanmaya Sethi were involved in the early stages of isolation of plant microtubules and Yash Jawale was involved in the early efforts of modeling tubulin polymerization kinetics. A grant from the Science and Engineering Research Board SERB, Govt. of India (EMR/2016/005481) to CAA supported the work. KJ was supported by a fellowship from DST-INSPIRE, Govt. of India (IF-130394) and a research associateship from SERB, Govt. of India (EMR/2016/005481). MR was supported by a fellowship from DST-INSPIRE, Govt. of India.

## Author contributions

Kunalika Jain: Investigation, methodology, formal analysis, reviewing & editing

Megha Roy: Methodology & formal analysis.

Chaitanya A. Athale: Conceptualization, funding acquisition, project administration, supervision, writing-original draft, writing-reviewing & editing.

## Supporting Information

### Supporting Tables

**Table S1:** Previously reported critical concentration of polymerization values for tubulin from multiple sources.

| **Tubulin**<br>**source** | **$c^*$** | **Reference** |
| --- | --- | --- |
| Rose cells | 1.9 $\mu\text{M}$ | (Morejohn and Fosket, 1984) |
| *Arabidopsis* | 7.6 $\mu\text{M}$ | (Hotta et al., 2016) |
| Lily pollen | 12 $\mu\text{M}$ | (XU et al., 2005) |
| Porcine brain | 5 $\mu\text{M}$ | (Walker et al., 1988) |
| Bovine brain | 5.6 $\mu\text{M}$ | (Davis et al., 1993) |
| *S. cerevisiae* | 1.8 $\mu\text{M}$ | (Davis et al., 1993, Bode et al., 2003) |

### Supporting Figures

**Figure S1:**
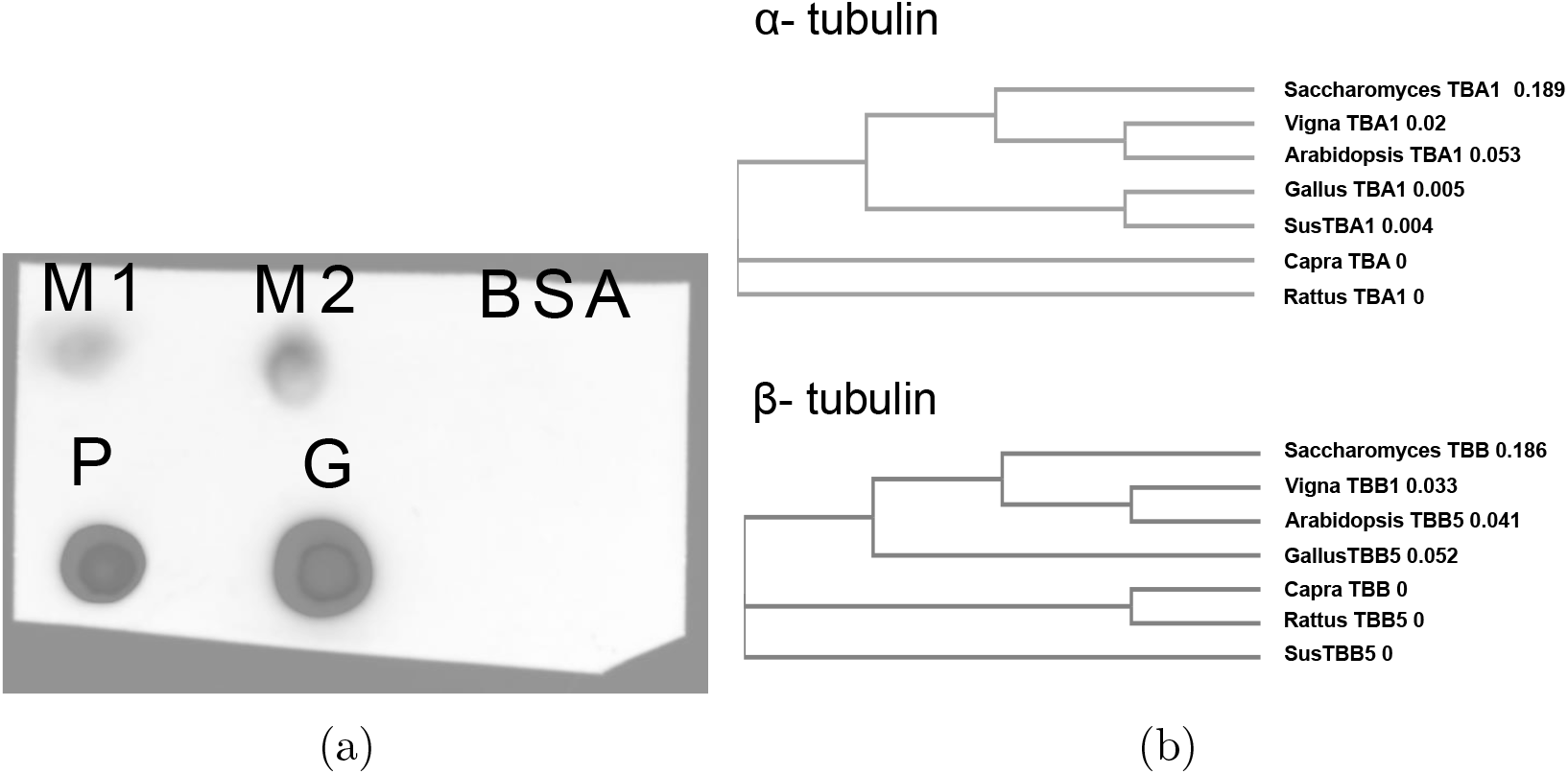
Dot blot and phylogenetic tree of related α- and β-tubulins. (a) Dot blots of mung tubulin samples (M1: 6 *µ*g and M2: 3.3 *µ*g, two independent isolates), porcine (P: 2 *µ*g) and goat brain tubulin (G: 1.26 *µ*g) and 2 *µ*g BSA were hybridized with a polyclonal anti-Arabidopsis *α*-tubulin antibody and detected by HRP conjugated anti-rabbit secondary antibodies and developer. (b) The phylogenetic tree based on protein sequence differences between the *(top) α*1- and *(bottom) β*-tubulins from *S. cerevisiae, Arabidopsis, Gallus* (chicken), *Rattus* (rat), *Sus* (pig) and *Capra* (goat) are compared to *Vigna* (mung bean). The tree was generated using Clustal Omega.

**Figure S2:**
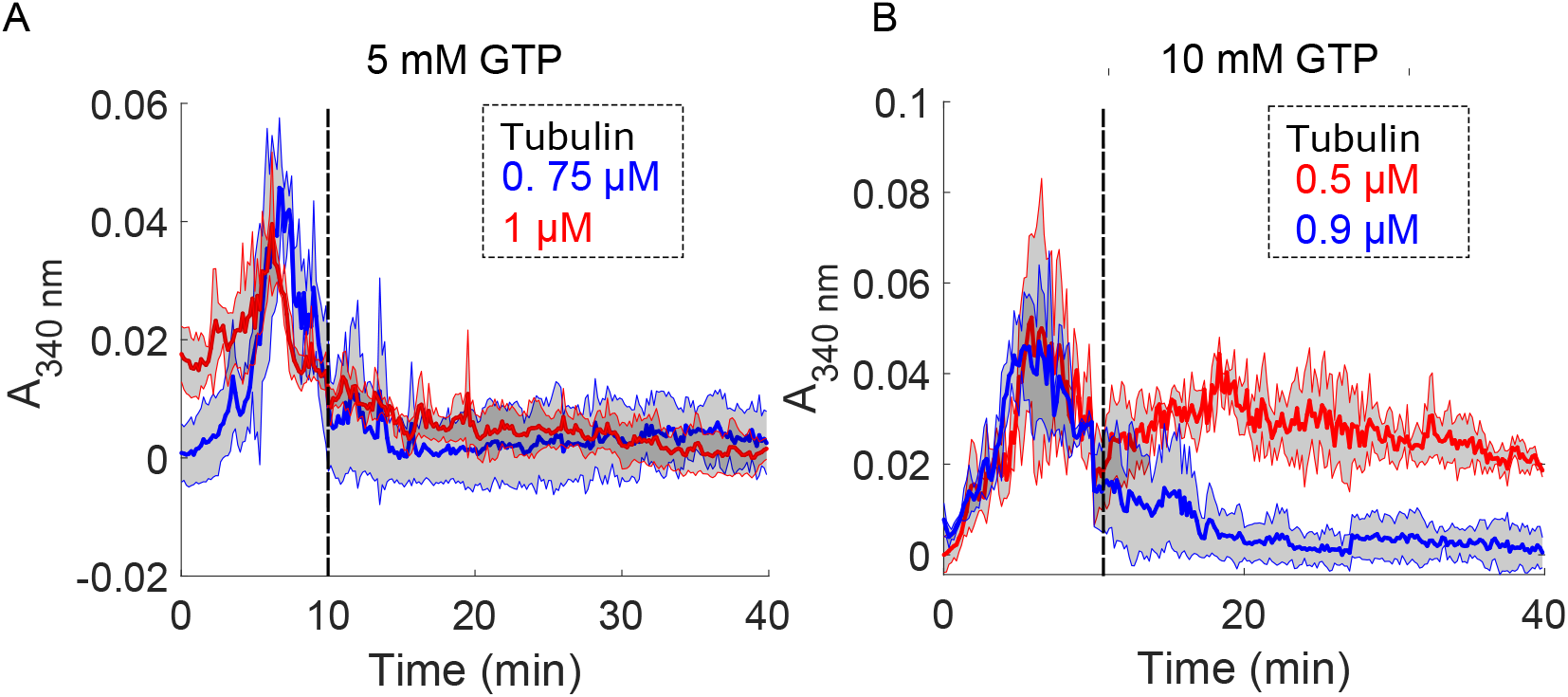
GTP hydrolysis as a limiting factor in mung tubulin polymerization kinetics. The polymerization kinetics of mung tubulin measured by the absorbance 340 nm over time with either (A) 5 mM or (B) 10 mM GTP added after 10 minutes (dashed vertical line). Tubulin concentration ranged from 0.5 to 1 *µ*M. Solid lines: mean data, colours: tubulin concentrations and grey area: ±s.d.

**Figure S3:**
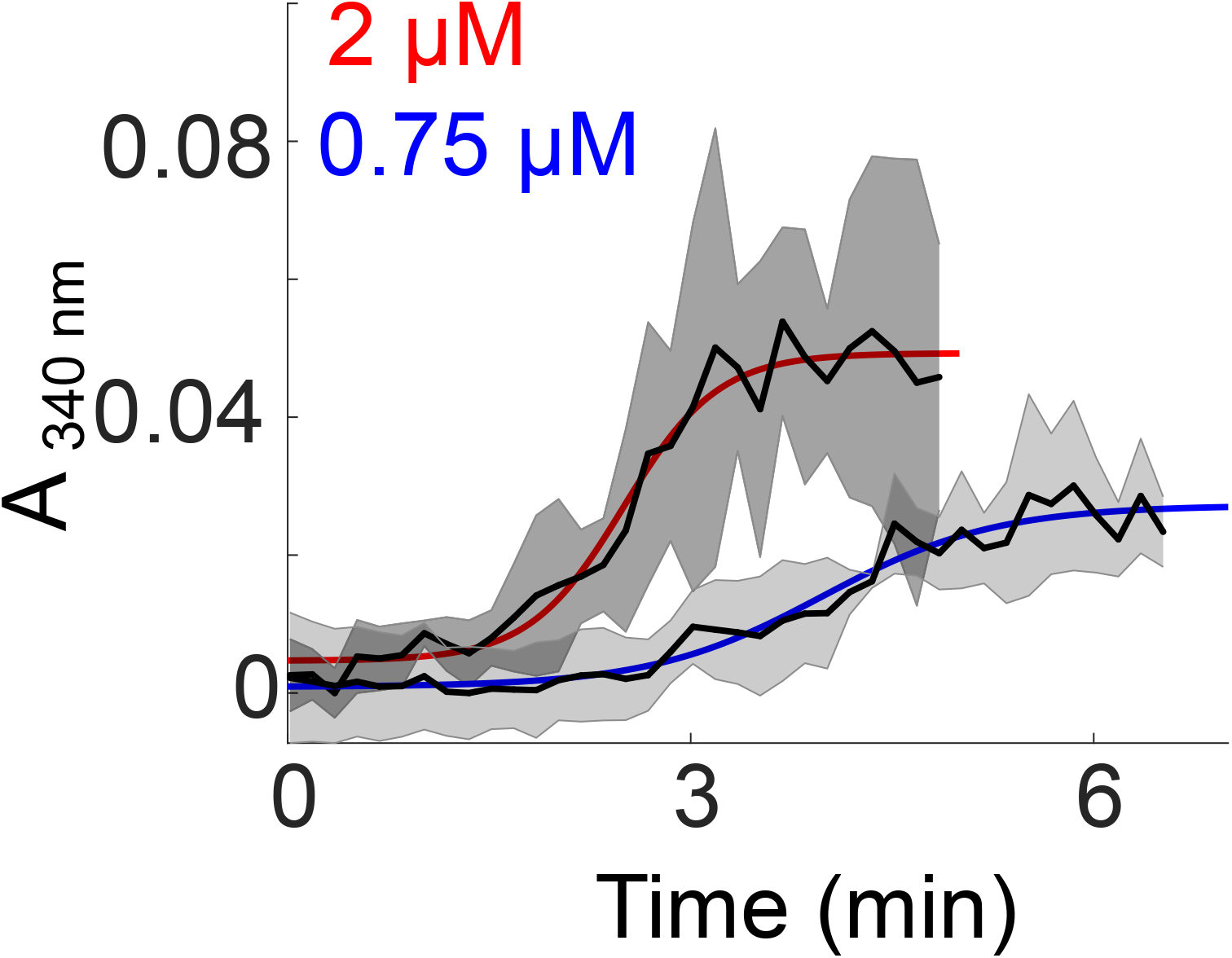
Taxol stabilized polymerization kinetics of mung tubulin. Polymerization kinetics of 0.75 and 2 *µ*M mung tubulin in presence of 10 *µ*M taxol were were measured by absorbance at 340 nm and the mean data (black line) ± s.d. (grey area) were fit to the saturation polymerization model (Equation 1).

### Supporting Video

**Video SV1:**
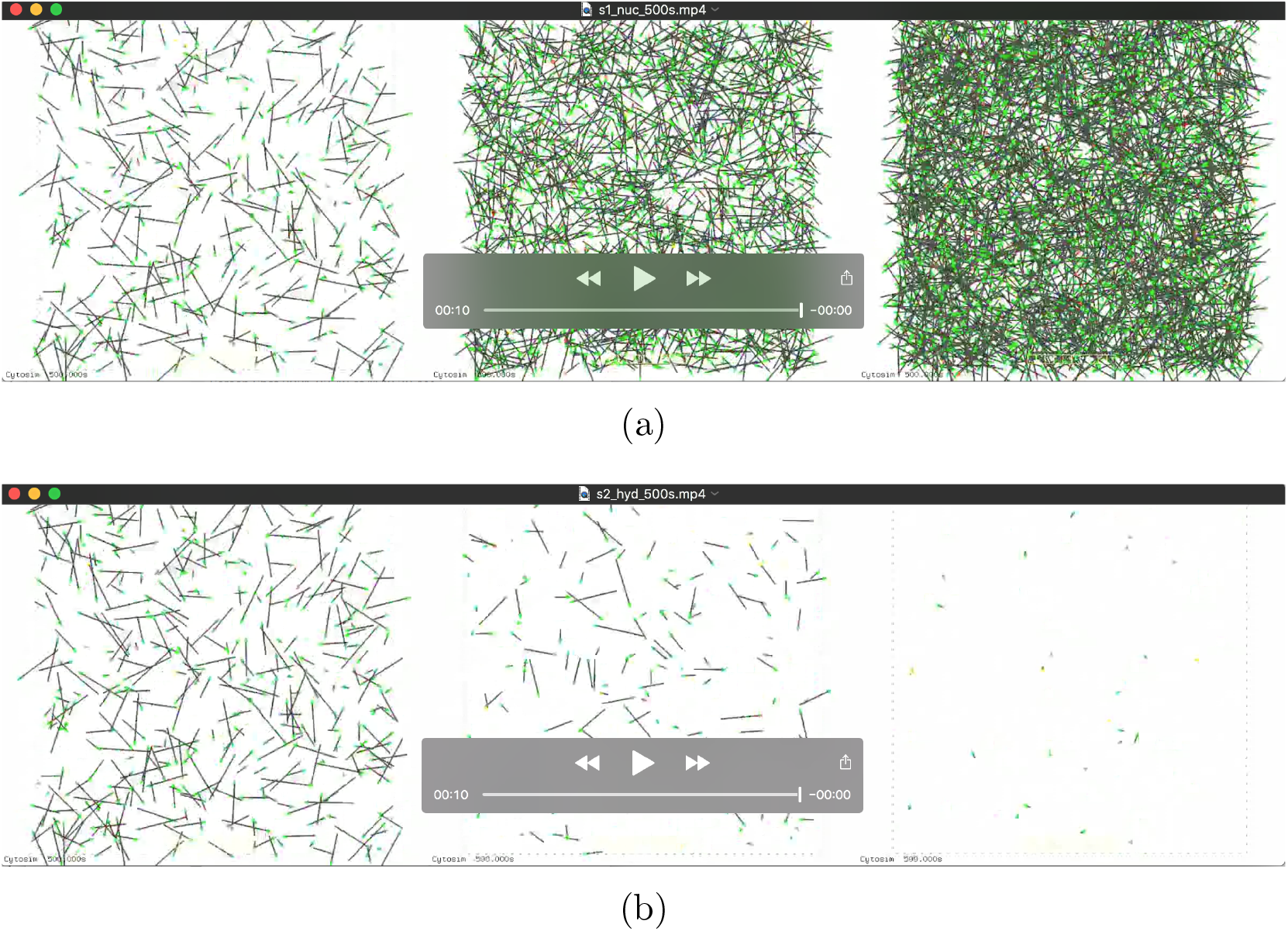
Effect of nucleation and hydrolysis rates on MT filament dynamics. MT growth was simulated as stochastic nucleation and elongation of multiple 1D filaments in a 2D square periodic box of size 40 *×* 40 *µ*m^2^ (black dotted line. Filaments were nucleated and polymerized where (a) the nucleation rate was increased Lelft *→* Right: 1 *s*^-1^, 5 *s*^-1^ and 10 *s*^-1^) while the hydrolysis rate was constant (0.02 *s*^-1^) or (b) the hydrolysis rates was varied Left *→* Right: 0.02 *s*^-1^, 0.06 *s*^-1^ and 0.18 *s*^-1^), while the nucleation rate was constant (1 *s*^-1^).

